# Multi-modal imaging reveals dynamic interactions of *Staphylococcus aureus* within human neutrophils

**DOI:** 10.1101/2021.03.05.434126

**Authors:** Yin Xin Ho, Elliot Steele, Lynne Prince, Ashley Cadby

## Abstract

*Staphylococcus aureus* is an important human pathogen that causes a wide range of infections. Neutrophils are an essential component of our innate immune system and understanding S. aureus-neutrophil interactions on a sub-cellular level is crucial to developing new therapeutic strategies to promote immunity during *S. aureus* infections. To this end we have developed a multi-modal imaging platform capable of following host-pathogen processes in biological systems, this is achieved by switching imaging modalities between a low photo-toxicity and low resolution imaging modality through an increasing illumination intensity to achieve live super-resolution imaging. This novel imaging platform was applied to the study of human neutrophils infected by *S. aureus*. We show that we can image different infection stages of *S. aureus* in live neutrophils with super resolution microscopy. We see evidence of binary fission occurring in intracellular *S. aureus* within a neutrophil.

## Introduction

*Staphylococcus aureus* is a highly opportunistic pathogen that can silently colonize the human nares and yet also cause a wide range of life-threatening infections, including skin and soft tissue infections, septicemia and osteomyelitis (1). The incidence and severity of *S. aureus* infections can be attributed to its ability to acquire and develop resistance toward antibiotics (2). Since the evolution of methicillin-resistant *S. aureus* (MRSA) strain, life-threatening MRSA infections have been an increasing burden in many countries, with particularly high prevalence in the United States and Asia (3). Furthermore, *S. aureus* has garnered major attention as a successful pathogen due to its extensive arsenal of virulence factors, including those that facilitate adhesion and colonisation through to evasion of host immune response (4, 5).

Neutrophils are professional phagocytes constituting the body’s first line of defence, and are critical in defending the host against *S. aureus* infections. Following neutrophil phagocytosis of *S. aureus*, the bacteria are contained within phagosomes, a membrane-bound compartment where most of the killing mechanisms occur. The phagosomal maturation process involves the fusion of neutrophil granules with the phagosome, which deploys proteases and antimicrobial peptides into the phagosomal space (6). NADPH oxidase and myeloperoxidase rapidly generate superoxide and HOCl which have strong antimicrobial properties (7, 8). Despite these antimicrobial strategies, neutrophils fail to eliminate internalised *S. aureus* completely (9–11). Visualising dynamic host-pathogen interactions on a individual subcellular level, as opposed to in fixed samples and on a cell population level, allows us to better understand how *S. aureus* circumvents the neutrophil response and ultimately reveal pathways to therapeutically target to promote immunity during *S. aureus* infections.

Among the challenges of following the sub-cellular processes of intracellular *S. aureus* within human neutrophils is the spatial resolution required to image the processes involved formation of a phagosome. The high resolution which can only be circumvented by super-resolution imaging techniques (12). However, unlike other cell types, there is little application of super-resolution live imaging to the study of human neutrophils. In general this is due to the high illumination power required to implement super-resolution modalities which induces photo-damage in cells. Neutrophils are notorious for their sensitivity to imaging conditions (13). For example, reactive oxidative species (ROS) production generated by high illumination power is toxic to cells, thus limiting the duration of live imaging (14). Therefore, low illumination power is required for imaging live neutrophils, and it is crucial in the context of neutrophil and *S. aureus* interaction studies, as ROS production is a key antimicrobial response against *S. aureus* (15). Therefore, we need to image neu-trophils with minimal illumination power to gain an accurate understanding of intracellular *S. aureus* dynamics within human neutrophils.

## Results

### Image Scanning Microscopy allows us to image the structure the phagosome membrane

The multi-modal platform used in this work allows us to visualise the host pathogen interactions of *S. aureus* and human neutrophils. An example of this is given in Figure 1a, and shows the time course of *S. aureus* and neutrophil interactions. The neutrophils were imaged for 80 minutes, where the first 60 mins were visualised using a low power multi-point confocal modality, and the final 20 minutes were visualised by switching to an Image Scanning Microscopy (ISM) modality. This provided higher resolution imaging at the cost of higher photo-damage. Additionally, image obtained using ISM modality have improved signal-to-noise ratio (SNR), as shown in Figure 1c. The added resolution gained by ISM also allows us to resolve the structure of the neutrophil phagosome membrane. For example, Figure 2 shows a subset of intracellular *S. aureus* present within neutrophil phagosomes (white arrows).

**Fig. 1.**
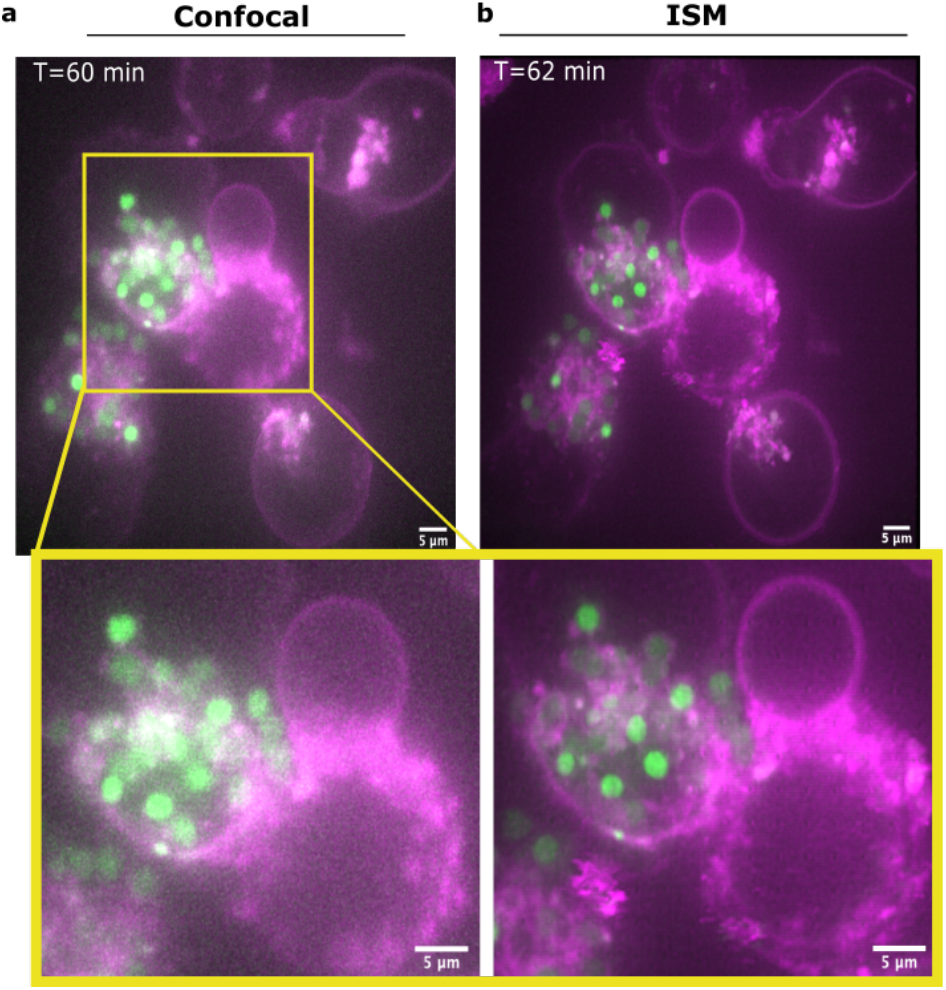
Multi-modal imaging. a) Time series of neutrophil and *S. aureus* visualised using multi-point confocal, followed by ISM. Images shown are time points taken from the a) 60 min (confocal) and b) 62 min (ISM), where neutrophil-*S. aureus* were imaged for 80 mins. An area (yellow box) is selected to highlight the differences in resolution and SNR between confocal (left panel) and ISM (right panel) modalities. Neutrophils were labelled with CellMask DeepRed plasma membrane stain, and coinfected with GFP-labelled *S. aureus* at multiplicity of infection (MOI) of 5. Images shown are z-projects displayed at maximum intensity projections.

**Fig. 2.**
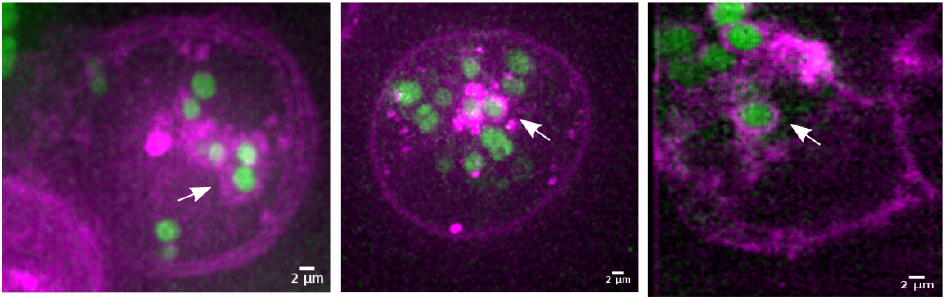
Live ISM imaging of neutrophil and *S. aureus*. Phagosome membrane encapsulating *S. aureus* (white arrows) can be observed within neutrophils across 3 independent experiments. Neutrophils were labelled with CellMask DeepRed plasma membrane stain, and co-infected with GFP-labelled *S. aureus* at multiplicity of infection (MOI) of 5. Images shown are z-projects displayed at maximum intensity projections.

### Heterogenous phagosomal acidification within neutrophils

This multi-modal approach to imaging gives us the ability to perform versatile multi-colour super-resolution imaging at specific time points allowing us to investigate the highly dynamic interactions between *S. aureus* and neutrophils with reduced photo-damage. Giving us the ability to follow processes at a lower spacial resolution with reduced photo damage, before switching to a higher resolution at the cost of higher photo damage. This combined with active dyes allows us to follow processes in phagocytosis. For example, dyes sensitive to the neutrophil micro-environment can be conjugated to GFP-labelled *S. aureus* to understand their dynamics within neutrophils. Here we used the pH sensitive pHrodo dye which allows us to visualise *S. aureus* within an acidified phagosome (16, 17). GFP-labelled *S. aureus* were stained with pHrodo red prior to co-infection with neutrophils. This enabled us to identify populations of *S. aureus* in acidified (magenta) and non-acidified (green) phagosomes within neutrophils. The early stages of *S. aureus* and neutrophil interactions were visualised in three colours using a multi-point confocal modality for 1 h, where a heterogenous population of *S. aureus* within acidified and non-acidified phagosomes were observed (Figure 3a). Interestingly, switching from confocal (Figure 3b) to higher resolution imaging of the samples by using ISM revealed cell wall structures only in the green (non-acidified) *S. aureus* (yellow arrows, Figure 3c). The cell wall structures in intracellular *S. aureus* were observed across 3 independent experiments (Supplementary Figure 5), and these structures were not visible in the confocal modality as shown in Figure 3b.

**Fig. 3.**
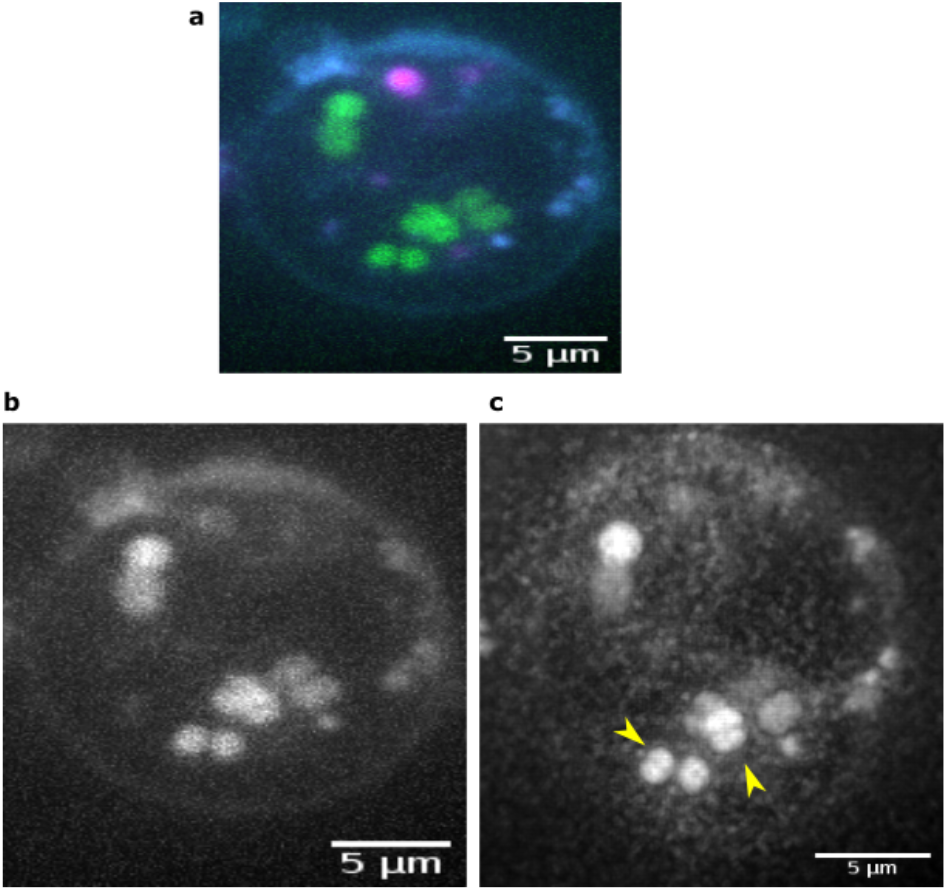
Confocal and ISM imaging of live neutrophils and *S. aureus*. a) Composite image of *S. aureus* and neutrophils visualised using a confocal modality. GFP-labelled *S. aureus* visualised using 470 nm laser line in b) confocal and c) ISM are compared to show the improvement in resolution using ISM. The yellow arrows showed the *S. aureus* with cell wall structures observed in ISM. The images shown are z-projects displayed at maximum intensity projections.

### Fluorescence D-amino acid labelling to visualise intracellular bacterial cell division

The formation of a cell wall implies that either a pair of *s. aureus* were phagocytosed mid-replications, or that cell division has occurred within in the neutrophil. To investigate this possibility we imaged *S. aureus* using dyes which specifically target new cell wall growth. Fluorescent D-amino acid dyes with intrinsic fluo-rophores attached, such as 7-hydroxycoumarin 3-carboxylic acid (HADA) enabled fast and real time detection of peptidoglycan cell wall synthesis in extensive range of bacterial species including *S. aureus* (18, 19). Here, for the first time, we apply this technique to visualise new cell wall growth within immune cells.

*In vivo* application of fluorescence D-amino acid was described previously (20). To ensure that the fluorescence D-amino acid labelling strategy used to probe the intracellular *S. aureus* dynamics does not affect neutrophil viability. We performed flow cytometry neutrophil cell death assay using ToPro-3 viability dye, and assessed the neutrophil morphology to determine the effect of fluorescence D-amino acid on neutrophil viability (Methods). Together, the flow cytometry and morphological viability assays confirmed that 1 mM of HADA used to label intracellular *S. aureus* does not compromise neutrophil viability (Supplementary Figures 7 and 8).To determine whether the cell wall structures observed in *S. aureus* indicated that intracellular *S. aureus* undergoes cell division within neutrophils, HADA was used to label intracellular live and heat-killed *S. aureus*. Figure 4 showed HADA labelling of intracellular live *S. aureus*, confirming the ability to apply fluorescence D-amino acid labels to probe intracellular *S. aureus* in live neutrophils. FDAA labelling were used to probe newly synthesised peptidoglycan cell walls of *S. aureus* (18, 19). Therefore, the incorporation of HADA into live intracellular *S. aureus* suggests that new cell wall is being synthesised, and potentially indicating early stages of replication within neutrophils.

**Fig. 4.**
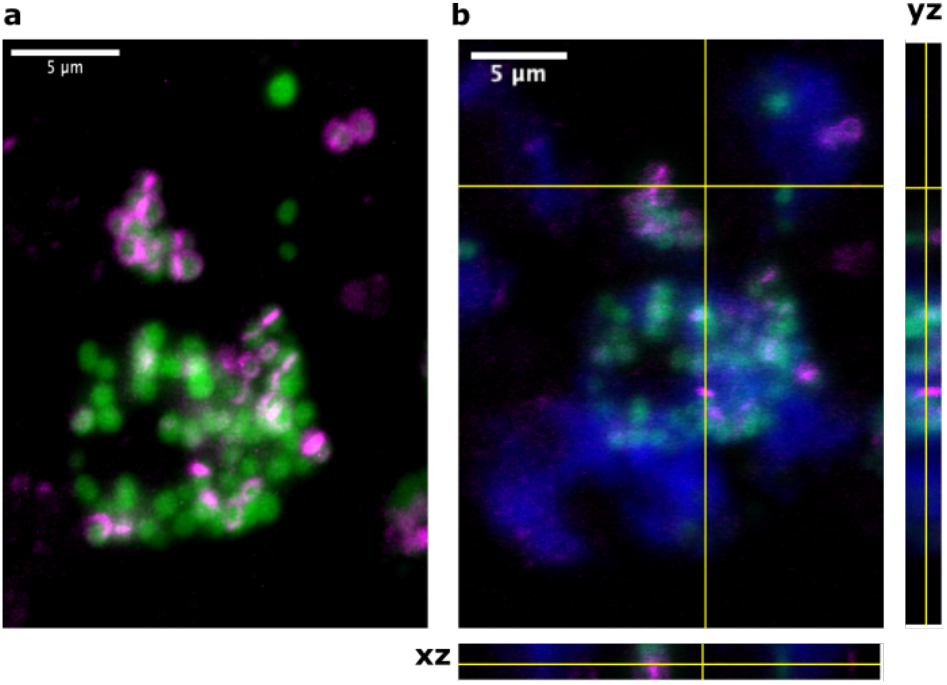
Fluorescence D-amino acid labelling of intracellular *S. aureus*. Neutrophils were co-infected with GFP-*S. aureus* at MOI of 5 for 2h, and 1mM of fluorescence D-amino acid label HADA was added to the co-culture 30 mins prior fixation to label intracellular *S. aureus*. a) HADA(magenta) labelling of intracellular *S. aureus* (green). The image is shown without the neutrophil membrane, with z-projects displayed at maximum intensity projections. b) Same image with neutrophil membrane (blue), displayed with orthogonal views to confirm the presence of HADA-labelled intracellular *S. aureus* within neutrophils.

## Discussion

Our implementation of a multi-modal imaging platform enabled super-resolution live imaging of human neutrophils, revealing the dynamics of intracellular *S. aureus* within neutrophils. Using GFP-*S. aureus* labelled with pHrodo red revealed the presence of *S. aureus* in acidified and non-acidified neutrophil compartments, suggesting that a population of live *S. aureus* were able to resist acidification in neutrophil phagosome. Imaging in high resolution using the SIM modality also enabled the visualisation of neutrophil phagosomal membrane encapsulating *S. aureus*. This further revealed a subset of *S. aureus* that were not encapsulated within the phagosomes in neutrophil. However, due to the reduced temporal limitation of the the SIM modality it was not possible to study the development of the phagosome membrane.

Using ISM, we also observed cell wall structures in *S. aureus* within non-acidified phagosomes. This suggests that *S. aureus* within non-acidified phagosomes undergo cell division within the neutrophils. To further investigate this, a novel application of fluorescence D-amino acid labelling approach in human neutrophils was used. Our data showed that fluorescence D-amino acid labels are permeable to neutrophil membrane, enabling their incorporation to live intracellular bacteria. Furthermore, we confirmed that the fluorescence D-amino acid labelling approach does not compromise neutrophil viabilty. We observed a subset of intracellular *S. aureus* labelled with HADA, which binds to newly synthesised peptidoglycan cell wall. The presence of HADA in intracellular *S. aureus* suggests that active peptidoglycan cell wall synthesis occurs within neutrophils. Additionally, various bacterial growth patterns, including peripheral and perpendicular labelling patterns in intracellular *S. aureus* were observed, thus indicating that cell wall synthesis occurs at various sites. This also suggests that the intracellular *S. aureus* undergo different growth phase within the neutrophils. However, due the spectral range of the Cairnfocal system, it was not possible to image HADA with a higher spatial resolution. However, the Cairnfocal did allow us to swap modalities such that HADA could be imaged in epi-fluorescence mode while all the other channels were imaged at high resolution. Given the versatility of the Cairnfocal system, it is possible to use a targeted approach to selectively visualise FDAA-labelled intracellular *S. aureus* in super-resolution modalities such as ISM or single-molecule localisation microscopy to further investigate the different bacterial growth patterns, as a way to reduce overall photo bleaching of samples.

Together, using super-resolution live imaging and biochemical labelling approaches we show that live intracellular *S. aureus* in non-acidified compartments of neutrophils are growing as shown by new peptidoglycan cell wall synthesis occuring in the *S. aureus* within non-acidified compartments.

## Materials and Methods

### Neutrophils isolation and culture

Human neutrophils were isolated by Plasma-Percoll density gradient centrifugation from whole blood of healthy donors based on previously described literature (21, 22). The study was carried out with written informed consent obtained from each donor. The ethical approval was obtained from the South Sheffield Research Ethics Committee (study number STH13927).

Freshly isolated neutrophils were resuspended at 5×10^6^ cells/ml in phenol red-free Roswell Park Memorial Institute (RPMI) 1640 media (Thermo Fisher, Waltham, MA), supplemented with 10% (v/v) heat-inactivated fetal bovine serum (FBS, PromoCell, Heidelberg, Germany) and 25mM HEPES buffer solution (Sigma Aldrich, Gillingham, Dorset). Neutrophils were cultured on high 35mm *μ*-dish with ibiTreat #1.5 polymer coverslip (ibidi, Martinsried, Germany).

### Bacterial culture and labelling

USA300 *S. aureus* strain JE2 with GFP chromosomal insertion were grown in Brain Heart Infusion (BHI) broth at 37°C with 5% CO_2_, with shaking at 350 revolutions per minute (rpm) until mid-log phase. The GFP-*S. aureus* were labelled with 2.5 mM pHrodo Red succinimidyl ester (Invitrogen, Carlsbad, CA) according to manufacturer’s instruction.

### Fluorescence D-amino acid labelling of intracellular *S. aureus*

Fluorescence D-amino acid labels were obtained from Dr. Victoria Lund and Professor Simon Foster (Department of Molecular Biology and Biotechnology, University of Sheffield) (19). Freshly isolated neutrophils (5×10^5^ cells) were stained with 10 *μ*g/ml of CellMask Deep Red plasma membrane stain (#C10046, Invitrogen, Carlsbad, CA), followed by co-infection with GFP-*S. aureus*, prepared as above, at multiplicity of infection (MOI) 5 for 30 mins. Extracellular bacteria were removed by centrifugation at 2000 rpm for 4 mins, and cells were resuspended in fresh phenol-red free RPMI (supplemented with 10% (v/v) heat-inactivated FBS and 25 mM HEPES). The cultures were left to incubate at 37°C with 5% CO_2_ until specified time points. To label the peptidoglycan of intracellular *S. aureus*, 1 mM HADA was added to the cultures 30 mins before specific time points. To remove excess dyes, cells were centrifuged at 2000 rpm for 3 mins, followed by washing with fresh phenol-red free RPMI 1640 (supplemented with 10% (v/v) FBS). To preserve cell structures for widefield imaging, the cells were plated on poly-L-lysine-coated coverslips and fixed with 4% paraformaldehyde (PFA) for 10 mins at room temperature, followed by washing twice with phosphate-buffered saline (PBS). Vectashield Antifade mounting medium (#H-1000, Vector Laboratories, Burlingame, CA) was added to the samples before mounting onto glass microscopic slides.

### Fixed cell imaging

Fixed samples were imaged using the inverted Zeiss LSM 510 NLO confocal microscope, with 63x/1.4 oil immersion lens. Excitation was delivered at 488 nm, 633 nm, 750 nm, and using FITC, CY5 and DAPI filters. The step sizes for z-stack acquisition was 633 μm.

### Neutrophil cell death assay

Neutrophil viability was assessed by flow cytometry using the Attune Autosampler (Thermo Fisher, Waltham, MA). Freshly isolated neutrophils (5×10^5^ cells) were cultured in RPMI 1640 media and HADA (0.5, 1 and 2 mM) at 37°C with 5% CO_2_ for 2h. ToPro-3 viability dye diluted to 1:10000 was added to the samples before flow cytometry. Forward scatter and side scatter profiles, as well as the ToPro-3 negativity of neutrophil media control were used to gate the viable neutrophils, which enabled the enumeration of absolute viable neutrophil and ToPro-3 negative neutrophil count. Flow cytometry analysis was performed using FlowJo software (TreeStar, Ashland, OR).

### Morphological assessment of neutrophil viability

Apoptotic neutrophils display condensed nuclei morphology. Neutrophils were cultured in RPMI 1640 media and 1 mM HADA respectively at 37^°^ C with 5% CO2 for 2h. Cytospin slides of neutrophil cultures were prepared by cytocentrifugation, followed by Diff-Quick (Thermo Fisher, Waltham, MA) staining. The cytospin slides were visualised using a light microscope, and at least 300 cells were scored in each experiment to assess the percentage of neutrophils with apoptotic morphological features.

### Live cell imaging

The live-cell Confocal and ISM were performed using the CairnFocal (Cairn Research, Kent, UK) based system described earlier and in SI 3. Excitation was delivered at 470 nm, 555 nm and 647 nm using the Laser Diode Illuminator (89-North, Vermont USA). The excitation light was filtered with a ZET405/470/555/640X quad-band excitation filter (Chroma Technology, Vermont USA). A matching dichroic mirror, the ZT405/470/555/640RPC (Chroma Technology, Vermont USA) was used to separate the excitation and emitted light. Emitted light was filtered via the ZET/405/470/555/640M emission filter before being detected by a Prime 95B sCMOS camera (Teledyne Photometrics, Arizona, USA). The system utilised an infinity corrected, 100x, 1.49NA, oil immersion TIRF lens, mounted on an Eclipse Ti microscope frame (Nikon Instruments, UK). A 1.5X C-Mount Fixed Focal Length Lens Extender (Edmund Optics, North Yorkshire, UK) was placed between the DMD and the camera to ensure that the 11 micron pixels would be small enough to exceed the Niquist criterion with a 100x lens. Z-stacks were performed by moving the sample using a Nano-Z100 z-stage (Mad City Labs, Wisconsin, USA). During imaging, cells were maintained at 37 °C using the UNO-T Stage-Top incubator (Okolab, Naples, Italy). CO_2_ levels were not controlled.

The camera data was collected via the PCIe interface, allowing for frame-rates of up to 80 fps in full frame. Unless otherwise stated, experiments were run using the center 50% of the chip, allowing frame rates of up to ~ 150 fps. Software control of the system was performed using the Micro-Manager 2.0-gamma software suite and custom software for control of the DMD. Acquisitions requiring more than XYCZT imaging (i.e., ISM) were performed using custom Micro-Manager beanshell scripts.

Single ISM data frames were reconstructed from acquisition of 100 confocal pattern frames acquired with a 10 ms integration time. Reconstruction was performed by identifying and localising confocal spots using the ThunderSTORM ImageJ plug-in (23), and subsequently performing the Sheppard summing algorithm, via a MATLAB script, to enhance resolution.

### Multi-modal imaging platform setup

Implementation of a multi-modal imaging platform to adapt according to the biological samples was presented previously (24). Briefly, the system is built around the CairnFocal, a Digital Micromirror Device (DMD) based confocal system from Cairn Research Ltd. (25). A DMD is placed at an image plane between the camera(s) and the microscope, as well as in the illumination pathway. Any pattern displayed on the DMD will therefore be projected onto the sample, as well as determining which of the two cameras (dubbed the “on” and “off” side cameras) the emitted light is directed to. By displaying a pattern similar to the pinholes of a spinning disk confocal microscope (SDCM), a confocal image will be formed on the on side camera due to the groups of on pixels acting as the pinholes of a spinning disk. This pattern can then be translated such that the entire sample is illuminated, either in a single camera frame to perform confocal or by synchronising the camera and DMD such that each frame displayed on the DMD is captured in a single camera exposure; this allows for a super-resolution image to be constructed via a Sheppard summing algorithm applied in post proccesing (26). Other microscopy techniques can be applied by displaying different patterns on the DMD. In this work we use a combination of far field epi-fluorescent, confocal microscopy and super resolution microscopy by pixel reassignment (26). For more information see SI 3.

### Image analysis

Image stacks were pre-processed in ImageJ. Maximum intensity Z-projections were used to represent the 3D stacks. The images were enhanced using automatic brightness and contrast settings. To segment individual bacteria, automatic threshold and watershed algorithm were applied to single channel images. The circularity of bacteria were measured using the built-in plugin on ImageJ (27), and custom-written Python code was used to trace bacterial circularity across time.

## Supporting information

Multimodal_Supplementary

## ACKNOWLEDGEMENTS

We would like to thank Simon Foster, Simon Jones and Victoria Lund for their help with acquiring and using the dye HADA. This work was funded by the Medical Research Council SHIELD consortium “Optimising Innate Host Defence to Combat Antimicrobial Resistance” MRNO2995X/1 YXH, LP and AC would like to acknowledge financial support from the SHIELD consortium. YXH is supported by a studentship from The Florey Institute, University of Sheffield. ES would like to thank Cairn Research and the DTP for financial support.

