## Supplementary material for "Multi-modal imaging reveals dynamic interactions of *Staphylococcus aureus* within human neutrophils": Multimodal_Supplementary

### Supplementary Note 1: Morphological variations between bacterial populations

In addition to septum-like structures, we observed morphological variation among bacterial populations using ISM. Green *S. aureus* exhibited ovoid morphology, whereas red *S. aureus* remained rounded (Figure 5). Therefore, the circularity of cell was used as a descriptor to quantify the morphological variation between bacterial populations. The bacterial circularity in ISM time series were measured over time (Methods), with heat-killed GFP-labelled *S. aureus* as a control. We analysed images of *S. aureus* co-infected with neutrophils from three independent donors (Supplementary Figure 6.) Live red phrodo positive *S. aureus* situated within an acidified phagosome have higher circularity values, thus a more rounded morphology than green *S. aureus* across the time series, and for each bacterial population, there were minimal changes in the circularity values along the time series. Furthermore, the circularity of heat-killed GFP-labelled *S. aureus* time series is greater than both live *S. aureus* populations (Figure 6).

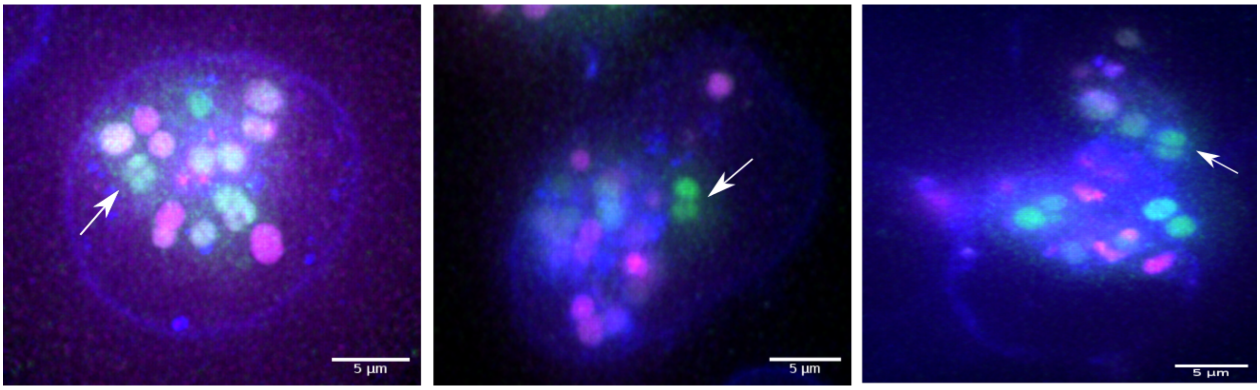


**Fig. 5. Visualisation of intracellular *S. aureus* in neutrophils.** Cell wall structures in intracellular *S. aureus* (white arrows) observed in 3 independent experiments by live imaging using ISM. Neutrophils were co-infected with *S. aureus* at an MOI 5, and the neutrophil membrane was labelled with Cellmask Deep Red (falsed coloured to blue), GFP-labelled *S. aureus* in green and pHrodo red in magenta. The images are z-projects displayed at maximum intensity projections.

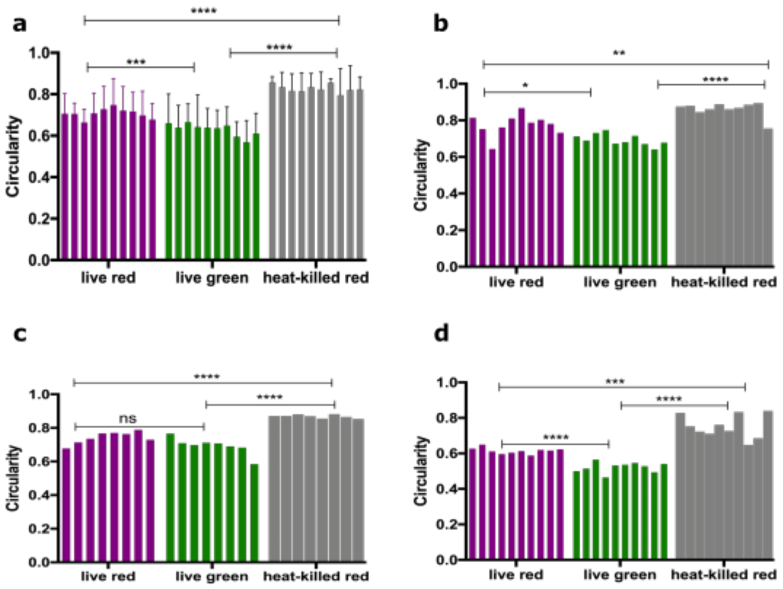


**Fig. 6. Comparison of the circularity between different bacterial populations.** The circularity of live red (in acidified compartment), green (non-acidified compartment) and heat-killed bacteria were measured and compared. Data shown in a) is the mean of 3 independent donors, and the measurements from independent donors b) donor 1, c) donor 2 and d) donor 3 were shown. Statistical analysis was performed using One-way ANOVA, **** p<0.0001,*** p<0.0002,* p<0.0332.

### Supplementary Note 2: Fluorescence D-amino acid labelling of intracellular *S. aureus*

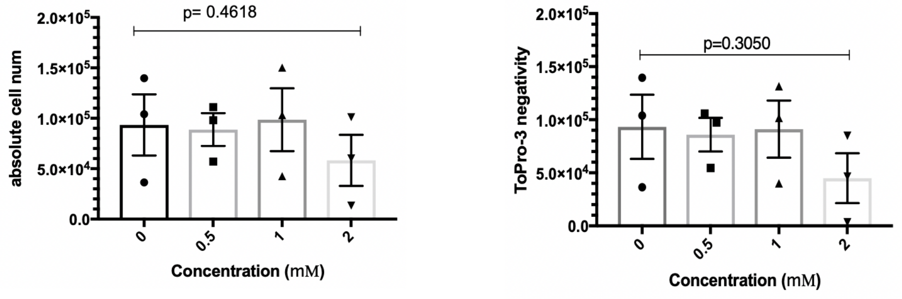


**Fig. 7. HADA does not affect neutrophil viability.** Different HADA concentrations (0.5, 1 and 2mM) were used in co-incubation with neutrophils for 2h, and neutrophil viability was quantified using flow cytometry, using ToPro-3 as neutrophil cell death marker. The absolute neutrophil cell number and ToPro-3 negativity decreased when 2mM HADA was used, indicating that 2mM of HADA is toxic to neutrophils. Data shown is the mean with standard error mean of 3 independent experiments. Statistical analysis was performed by One-way ANOVA.

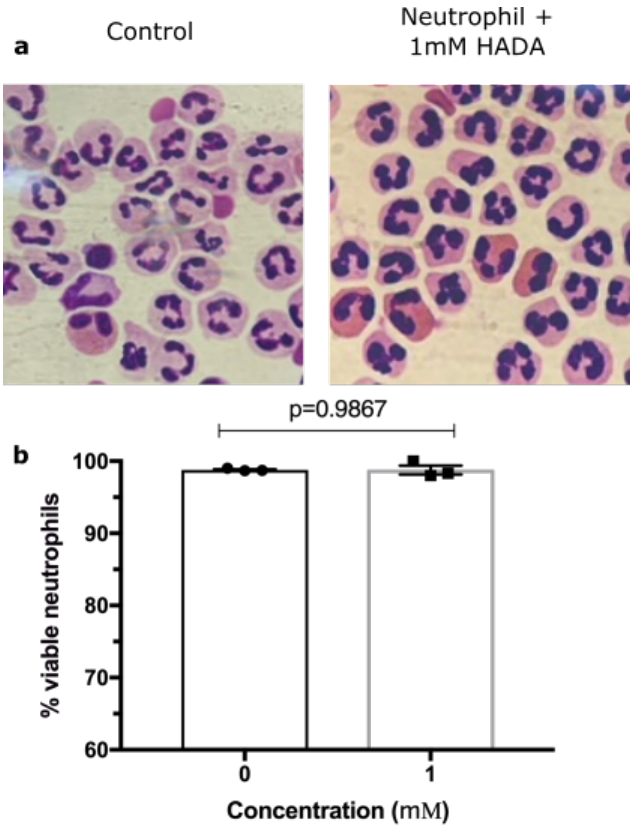


**Fig. 8. Neutrophil apoptosis assay.** a) Morphological assessment of neutrophils cultured in media as the control and 1mM HADA for 2h using light microscopy. b) The percentage of viable neutrophils between media control and 1 mM HADA treatment. Data shown is the mean with standard error mean of 3 independent experiments, and statistical analysis was performed by paired t-test.

### Supplementary Note 3: Imaging System

The multi-modal system described in this work is built around the CairnFocal DMD based confocal (Cairn Research, Kent, UK). Excitation was delivered at $470\,\mathrm{nm}$, $555\,\mathrm{nm}$ and $647\,\mathrm{nm}$ using the Laser Diode Illuminator (89-North, Vermont USA). The excitation light was filtered with a ZET405/470/555/640X quad-band excitation filter (Chroma Technology, Vermont USA). A matching dichroic mirror, the ZT405/470/555/640RPC (Chroma Technology, Vermont USA) was used to separate the excitation and emitted light. Emitted light was filtered via the ZET/405/470/555/640M emission filter before being detected by a Prime 95B sCMOS camera (Teledyne Photometrics, Arizona, USA). The system utilised an infinity corrected, 100x, 1.49NA, oil immersion TIRF lens, mounted on an Eclipse Ti microscope frame (Nikon Instruments, UK). A 1.5X C-Mount Fixed Focal Length Lens Extender (Edmund Optics, North Yorkshire, UK) was placed between the DMD and the camera to ensure that the 11 micron pixels would be small enough to exceed the Niquist criterion with a 100x lens. Z-stacks were performed by moving the sample using a Nano-Z100 z-stage (Mad City Labs, Wisconsin, USA). During imaging, cells were maintained at $37\,°\mathrm{C}$ using the UNO-T Stage-Top incubator (Okolab, Naples, Italy). $CO_2$ levels were not controlled.

**The CairnFocal.** The CairnFocal places the DMD in a plane conjugate with the sample in both the excitation and emission pathway. Excitation light is focused onto the DMD via 4 mirrors (2 flat mirrors, 1 concave and 1 convex arranged in a Schwarzschild relay configuration) (25). If the excitation light strikes a mirror in the 'on' position it is directed towards the back of the objective via a pair of relay lenses and the microscope tube lens. If the light strikes an 'off' mirror then the light does not reach the sample. As the DMD is placed in a conjugate plane, any image displayed on the DMD will be focused onto the sample. Illuminating via the CairnFocal's second illumination port works in an identical manner, however, flips which mirror positions are considered 'on' and 'off'.

Emitted light captured by the objective will be focused onto the DMD by the same optics used for excitation. Upon striking the DMD, the emitted light will be directed, via a Schwarzschild relay, to one of two cameras depending on whether it hits a mirror in the 'on' or 'off' position. The Schwarzschild relays act to correct out the $24°$ shift introduced by using the DMD in reverse, which would otherwise result in extreme distortions at the detection plane (25).

**DMD Patterns.** With the CairnFocal, multi-modality is achieved by changing the images displayed on the DMD. Widefield is achieved by placing all the mirrors in the 'on' position. This causes all light striking the DMD to be directed towards the sample, and all emitted light to be directed to the 'on' side camera (24)

Confocal operation is achieved by turning on a small group of mirrors surrounded by 'off' mirrors (24). A sufficiently small group of mirrors will be imaged onto a region smaller than the diffraction limit of the microscope in the sample plane. Light emitted from this region will be focused back onto the same small group of mirrors which will then act as a pinhole in a traditional confocal setup; in focus light will hit these 'on' mirrors and be directed to one camera, while out of focus light will strike the surrounding 'off' mirrors and be directed to the other camera. This group of mirrors can then be moved so as to sweep the entire field of view, building up an optically sectioned image on the 'on' camera and an image consisting of the rejected light on the 'off' camera. Spinning disk confocal imaging (SDC) is emulated by using multiple groups of 'on' mirrors simultaneously. An interesting advantage of this method over traditional SDC is that it is possible to change the spacing between pinholes and even the size of the pinholes during imaging. Increasing the spacing between pinholes reduces crosstalk between pinholes and improves optical sectioning at the expense of imaging speed. Increasing pinhole size results in worsened optical sectioning but increased signal to noise ratio. In this work a pinhole size of 3x3 mirrors ($41.1\,\mathrm{\mu m}$) and a pinhole spacing of 7 mirrors ($95.9\,\mathrm{\mu m}$) was found to provide a good balance of optical sectioning, imaging speed, SNR and phototoxicity. In principal the virtual pinholes displayed on the DMD could take several shapes, however, in practice the difference between shapes made of so few mirrors is negligible and as a result square groups were used for ease of implementation.

In confocal operation, the DMD is synchronised with the camera exposure such that the DMD displays the entire confocal pattern an integer number of times in a single exposure (e.g., for an exposure time of $100\,\mathrm{ms}$ and a confocal pattern requiring 100 frames to cover the entire field, each frame could be shown for $1\,\mathrm{ms}$, $0.5\,\mathrm{ms}$, $0.25\,\mathrm{ms}$, etc., but not $0.3\,\mathrm{ms}$ as this would result in 330 images being displayed in a single exposure and an inconsistent exposure time throughout the field of view). Alternatively, the camera and DMD can be synchronised such that a single camera exposure is performed per DMD frame. Each image will therefore contain a set of point spread functions (PSF) from pinholes that have not moved during that frame's exposure. Summing these frames will recover the standard confocal images, albeit at a significant read noise penalty when compared to the single exposure confocal described above. The images from this approach, however, contain extra information due to having access to each individual PSF. These PSFs can be processed via a Shepard Summing algorithm to provide an $\sim 1.4\times$ increase in resolution followed by an optional deconvolution step to provide a total increase in resolution of $2\times$.

**ISM Reconstruction Algorithm.** In this work, ISM reconstruction was performed via a combination of freely available ImageJ plugins and custom MATLAB scripts. Firstly, the location of the PSFs in each frame was located using the ThunderSTORM ImageJ plugin (23). These localisations were exported as CSV and loaded along with the original image data into custom MATLAB scripts which performed Shepard Summing (26) as follows:

1. An output image of width and height $8\times$ that of the input images is created and all values are set to $0$
2. A region around each PSF is cropped from the original image and upscaled by a factor of $4\times$
3. Each PSF is then added to the output image at the position corresponding to its centre in the input image
4. Once all PSFs have been incorporated into the output image, it is downscaled by a factor of $4\times$ and filtered with a low pass filter with a cut-off frequency greater than $2\times$ the diffraction limit to help remove reconstruction artifacts

The optional deconvolution step was not performed in this work.
